# A Model for Brain Life History Evolution

**DOI:** 10.1101/050534

**Authors:** Mauricio González-Forero, Timm Faulwasser, Laurent Lehmann

## Abstract

Mathematical modeling of brain evolution is scarce, possibly due in part to the difficulty of describing how brain relates to fitness. Yet such modeling is needed to formalize verbal arguments and deepen our understanding of brain evolution. To address this issue, we combine elements of life history and metabolic theories to formulate a metabolically explicit mathematical model for brain life history evolution. We assume that some of the brain’s energetic expense is due to production (learning) and maintenance (memory) of skills (or cognitive abilities, knowledge, information, etc.). We also assume that individuals use skills to extract energy from the environment, and can allocate this energy to grow and maintain the body, including brain and reproductive tissues. Our model can be used to ask what fraction of growth energy should be allocated to the growth of brain and other tissues at each age under various biological settings as a result of natural selection. We apply the model to find uninvadable allocation strategies under a “me-against-nature” setting, namely when overcoming environmentally determined energy-extraction challenges does not involve any interactions with other individuals (possibly except caregivers), and using parameter values for modern humans. The uninvadable strategies yield predictions for brain and body mass throughout ontogeny, as well as for the ages at maturity, adulthood, and brain growth arrest. We find that (1) a me-against-nature setting is enough to generate adult brain and body mass of ancient human scale, (2) large brains are favored by intermediately challenging environments, moderately effective skills, and metabolically expensive memory, and (3) adult skill number is proportional to brain mass when metabolic costs of memory saturate the brain metabolic rate allocated to skills. Overall, our model is a step towards a quantitative theory of brain life history evolution yielding testable quantitative predictions as ecological, demographic, and social factors vary.

**Author Summary:** Understanding what promotes the evolution of a given feature is often helped by mathematical modeling. However, mathematical modeling of brain evolution has remained scarce, possibly because of difficulties describing mathematically how the brain relates to reproductive success, which is the currency of evolution. Here we combine elements of two research fields that have previously been successful at detailing how a feature impacts reproductive success (life history theory) and at predicting the individual’s body mass throughout its life without the need to describe in detail the inner workings of the body (metabolic theory). We apply the model to a setting where individuals must extract energy from the environment without interacting with other individuals except caregivers (“me-against-nature”) and parameterize the model with data from humans. In this setting, the model can correctly predict a variety of human features, including large brain sizes. Our model can be used to obtain testable quantitative predictions in terms of brain mass throughout an individual’s life from assumed hypotheses promoting brain evolution, such as harsh environments or plentiful social interactions.

## Introduction

Empirical brain evolution research makes extensive use of correlational analyses seeking to test verbal hypotheses. For instance, correlations between diet quality or group size with cognitive ability or proxies thereof are routinely used to assess verbal hypotheses for whether ecological and social factors favor enhanced cognition or relatively large brains [1–8]. Functional studies have also been used to gain insight regarding brain evolution. For example, behavioral experiments report refined cognitive skills for social rather than general function [9,10], and brain imaging has identified various brain regions specialized for social interaction [11,12]. More recently, studies have addressed more directly the causes of large-brain evolution via phylogenetic analyses, artificial selection experiments, and genomic patterns of selection [13–17]. However, in contrast to other areas of evolutionary research, mathematical theory offering causal understanding and yielding testable hypotheses for brain evolution is still rare [18–20].

Mathematical modeling of brain evolution faces the trade-off between describing how brain impacts fitness without being overwhelmed by brain mechanistic details and at the same time considering enough mechanistic details to be able to make testable predictions. Existing models have described brain’s impact on fitness as facilitating energy acquisition from the environment allowing this energy to be used to increase survival [21], as facilitating energy production and/or decreasing the probability of being scrounged by others [22], as increasing offspring survival via parental care despite increasing mortality at birth [23], as increasing collaborative efficiency [24], as increasing mating ability [25], and as increasing the complexity of decision making regarding cooperation [26]. While these models have contributed to the qualitative understanding of brain evolution, evolutionary mathematical models yielding quantitative predictions, such as ontogenetic patterns for brain and body size, are still lacking.

Life history theory has been successful at making predictions about trait evolution by describing trait’s effects on fitness through explicit consideration of how an organism allocates its energy throughout ontogeny [27–31]. In turn, metabolic theory has been successful at making quantitative predictions about ontogenetic body mass and its focus on metabolism allows using a top-down perspective without the need to describe the inner functioning of the system [32–36]. Here, we combine elements of these two approaches to derive a model for brain life history evolution. This merging allows determining an individual’s optimal strategies regarding its ontogenetic energy allocation to the growth of its different tissues while obtaining quantitative predictions for body and brain size under different biological settings.

In linking brain to fitness, our model builds on previous models considering brain as embodied capital invested in fitness [21]. We consider separately the physical and functional embodied capital, the former being brain itself and the latter being skills (or cognitive abilities, knowledge, information, etc.) generated by the brain during ontogeny. Our consideration of ontogenetic skill accounts for the notion that information gained and maintained by the brain during ontogeny should be explicitly considered when attempting to understand brain evolution [21,37–40]. Thus, a defining feature of our model is that it assumes that some of the brain’s energetic consumption is due to acquisition and maintenance of skills (or cognitive abilities, knowledge, information, etc.). In turn, we consider skills that allow overcoming energy extraction challenges the individual faces at each age, and in so doing the individual obtains some energetic reward. The model is general in that various biological settings can be studied depending on the type of challenge individuals face at each age and on who engages in overcoming the challenge: the challenge can be ecological if it is posed by the non-social environment, or it can be social if it is posed by social partners; also, the individual can engage in overcoming the challenge either alone or in concert with social partners [these settings can be thought of as me (or us) against nature (or them)] [21,24,38,40–42].

We apply our model to analyze the baseline setting where individuals face exclusively ecological (non-social) challenges which are overcome by the individual alone (“me-against-nature” [24]). Then, given that the brain consumes some of its energy to gain and maintain skills and given the various types of challenges that the individual faces at each age, the model allows to predict how much an individual should grow its brain to obtain the energetic returns from skills. By feeding the model with parameter values for modern humans, we show how the model can yield predictions for life history stages as well as ontogenetic body and brain mass.

## Model

### Biological scenario

We consider a randomly mating population of large and constant size, where the environment is constant, generations are overlapping, individuals’ age is measured in continuous time, and the focus is on female survival and reproduction (i.e., standard demographic assumptions for life history evolution [28,31,43–46]). We partition the body of each female into three types of tissues (or cells): reproductive tissue, brain tissue, and the remainder tissue, which we refer to as somatic. To have energy at each age for body growth, body maintenance, and reproduction, each female extracts energy from its environment (e.g., by locating food, or by making resources usable through cracking or cooking), possibly with the help of her parents or caregivers (parental or alloparental care) and/or by interacting with other individuals in the population (e.g., through cooperative gathering or social competition for resources). To extract energy, each individual is assumed to use a number of relevant energy-extraction skills, which are produced and maintained by the brain.

We aim to determine the optimal allocation strategy of an individual’s energy budget to the growth of the different tissues throughout its lifespan, which is a form of the central life history question [28,31,43–46]. An allocation strategy is here a vector of evolving traits that is a function of the individual’s age, and that determines the individual’s energy allocation to the growth of its different tissues throughout the individual’s lifespan. To analyze how selection affects the evolution of the allocation strategy, we carry out an evolutionary invasion analysis (e.g., [31,47–49]), and thus consider that only two strategies can occur in the population, a mutant u and a resident (wild-type) 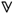 allocation strategies. We thus seek to establish which strategy is resistant to invasion by any alternative strategy taken from the set 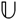 of feasible allocation strategies, and which thus provides a likely final point of evolution. From demographic assumptions we make below, it is well established [43, 50–52] that an uninvadable strategy 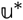 satisfies

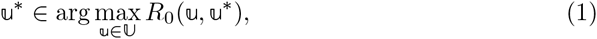

which implies that 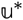 is a best response to itself, where

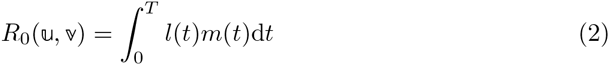

is the basic reproductive number of a single mutant in an otherwise monomorphic resident population and *T* is an age after which the individual no longer reproduces or is dead. The basic reproductive number depends on the probability *l*(*t*) that a mutant individual survives from birth until age *t* and on its rate *m*(*t*) of offspring production at age *t* with density dependence (“effective fecundity” [53], or the expected number of offspring produced at age *t* per unit time with density dependence), where these two vital rates may be functions of mutant and resident traits, 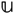 and 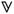.

To determine the lifetime offspring production R_0_ and how it connects to the state variables (tissues and skill) and to the evolving traits, we relate brain and skill growth to vital rates, which in turn is mediated by the connection between energy extraction, metabolism, and tissue growth. We thus formally derive our model by making these connections.

### Tracking resting metabolic rate

Standard life history models refer to complete components of the energy budget (e.g., assimilated energy; [54]). In practice, it is easier to measure heat release (metabolic rates; [55]). Hence, to facilitate empirical parameter estimation, we follow the approach of [34] and formulate our life history model in terms of resting metabolic rate allocation, rather than energy budget allocation. Thus, we track how resting metabolic rate is due to growth and maintenance of different tissues, in particular the brain.

We start from the partition of the individual’s energy budget used by [35], which divides the energy budget (assimilation rate) into heat released at rest (resting metabolic rate) and the remainder (Fig. 1; see [55] for details into why this partition is correct). The amount of energy used per unit time by an individual is its assimilation rate. Part of this energy per unit time is stored in the body (*S*) and the rest is the total metabolic rate, which is the energy released as heat per unit time after use. Part of the total metabolic rate is the resting metabolic rate *B*_rest_ and the remainder is the energy released as heat per unit time due to activity *B*_act_. In turn, part of the resting metabolic rate is due to maintenance of existing biomass *B*_maint_, and the remainder is due to production of new biomass *B*_syn_. We refer to *B*_syn_ as the growth metabolic rate (Fig. 1). We formulate our model in terms of allocation of growth metabolic rate *B*_syn_ to the growth of the different tissues.

**Fig. 1.**
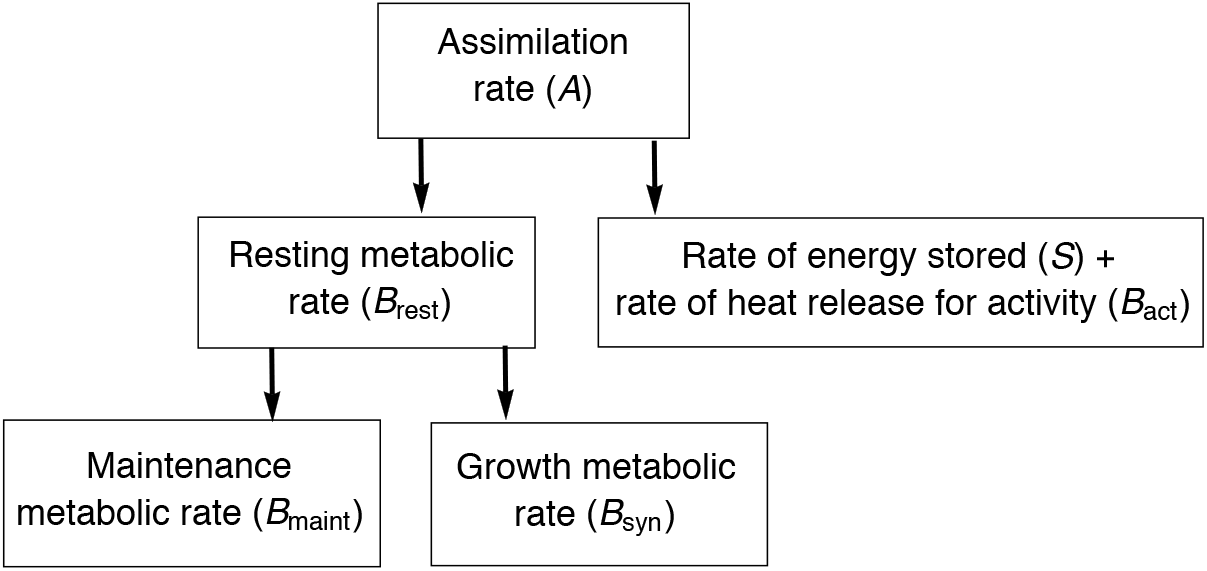
Partition of energy budget. Note the relation of resting metabolic rate to assimilation rate. Modified from [35].

### Energy partitioning

Denote by *N_i_*(*t*) the number of cells of type *i* of a focal mutant female of age *t*, where *i* ∈ {*b, r, s*} corresponds to brain, reproductive, and the remainder cells which we refer to as somatic, respectively. Assume that an average cell of type *i* in the resting body releases an amount of heat *B_ci_* per unit time. Hence, the total amount of heat released per unit time by existing cells in the resting individual is

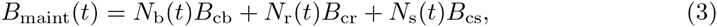

which gives the part of resting metabolic rate due to body mass maintenance [35].

Denote by 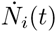 the time derivative of *N_i_(t)*. Assume that producing a new average cell of type *i* releases an amount of heat *E_ci_*. Hence, the total amount of heat released per unit time by the resting individual due to production of new cells is

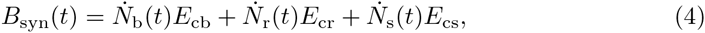

which gives the rate of heat release in biosynthesis [35], and we call it the growth metabolic rate. From (4), we have that

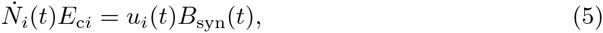

for *i* ∈ {*b, r, s*}, where *u_i_(t)* is the fraction of the growth metabolic rate due to production of new type-*i* cells at time *t* [summing over all cell types in (5) returns (4)]. The resulting time sequence 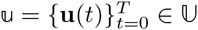, where **u**(*t*) = (*u*_b_(*t*), *u*_r_(*t*), *u*_s_(*t*)), of allocations from birth to (reproductive) death is the evolving multidimensional trait in our model and 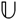 is the set of all feasible allocations strategies.

From our partitioning in Fig. 1, the total amount of heat released by the resting individual per unit time at age *t*

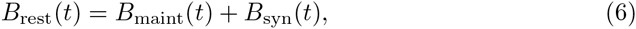

which is the individual’s resting metabolic rate at age *t*.

### Tissue growth rate

Let the mass of an average cell of type *i* be *m_ci_* for *i* ∈ {*b, r, s*}. Then, the mass of tissue *i* at age *t* is

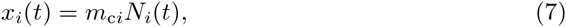

and hence, using (5), we have that the growth rate in mass of tissue *i* is

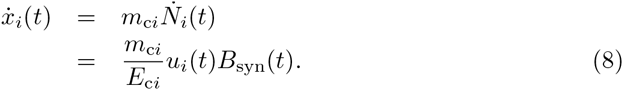

Denoting the heat released for producing a mass unit of tissue *i* as *E_i_* = *E_ci_*/*m_ci_*, this gives

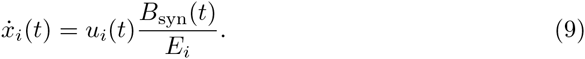

Using (6) in (9), we obtain the model’s first key equation specifying the growth rate of tissue *i*:

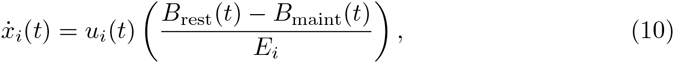

where from (3), we have that

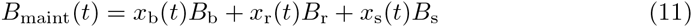

and the mass-specific cost of tissue maintenance is *B_i_* = *B_ci_*/*m_ci_*.

### Skill learning rate

We assume that some of the brain metabolic rate is due to acquiring and maintaining energy-extraction skills. We assume that the individual at age *t* has a number *x_k_*(*t*) of skills that can be used to overcome challenges of energy extraction. Denote by *M*_brain_(*t*) the brain metabolic rate of the individual at age *t* (i.e., the heat released by the brain per unit time with the individual at rest). From energy conservation, the brain metabolic rate must equal the heat released by the brain per unit time due to brain growth and brain maintenance; that is, from (3) and (4), the brain metabolic rate must satisfy

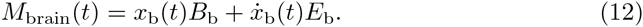

Let *s_k_* be the fraction of brain metabolic rate allocated to energy extraction skills, which we assume constant for simplicity. Suppose that the brain releases an amount of heat *E_k_* for acquiring an average energy-extraction skill (learning cost). Similarly, assume that the brain releases an amount of heat *B_k_* per unit time for maintaining an average energy-extraction skill (memory cost). Hence, from energy conservation, the rate of heat release by the brain due to skill growth and maintenance must equal the brain metabolic rate due to energy-extraction skills:

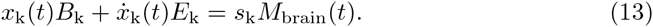

Rearranging, we obtain the model’s second key equation specifying skill learning rate:

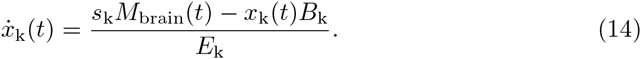

In analogy with (10), the first term in the numerator of (14) gives the heat released due to energetic input for learning whereas the second term gives the heat released for memory. [Note that an equation for skill growth rate can be similarly derived, not in terms of allocation to skill growth *and* maintenance *s_k_*, but in terms of allocation to skill growth u_k_ as for (10).]

### How skill affects energy extraction

We now derive an expression that specifies how brain affects energy extraction in the model. We consider that energy extraction depends on the focal female’s skills but possibly also on the skills of other females in the population. To make this dependence explicit, we denote by 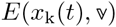 the amount of energy extracted by the focal female at time *t* from the environment, which depends on the individual’s skill *x_k_*(*t*) (and possibly body mass) and also on the skill or other features (state or control variables) of the resident population which ultimately depend on the resident allocation strategy 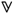 (so 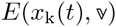) is assimilated energy plus surplus). Let *E*_max_(*t*) be the amount of energy that the individual obtains from the environment per unit time at age *t* if it is maximally successful at energy extraction (which also possibly depends on body mass). We define the energy extraction efficiency 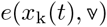 at age *t* as the normalized energy production per unit time at age *t*:

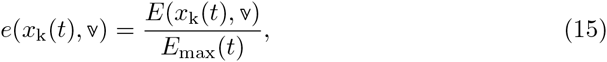

which is thus a dimensionless energy extraction performance measure.

We also define the ratio of resting metabolic rate to energy obtained per unit time as

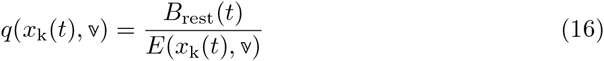

and, motivated by (16), we define

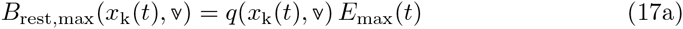

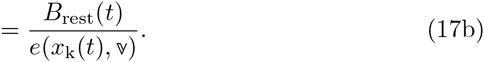

From (17b), we have that

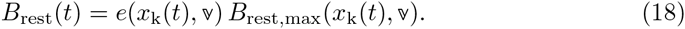

Consequently, 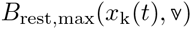 gives the resting metabolic rate when the individual is maximally successful at energy extraction.

Adult resting metabolic rate typically scales with adult body mass as a power law across all living systems [56–59], and also ontogenetically in humans to a good approximation (Fig. S3; but see [60]). We assume that this scaling holds for maximally successful individuals at energy extraction (assuming the scaling is empirically obtained from measurements in mostly well-fed individuals); that is, we assume

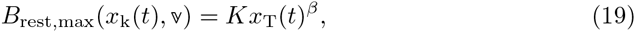

where *β* is a scaling coefficient and *K* is a constant independent of body mass (while both possibly depend on the resident strategy; note that *β* need not be 3/4). We further assume that energy extraction efficiency 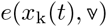 is independent of body mass, whereby Eqs. (18) and (19) yield the model’s third key equation specifying resting metabolic rate as:

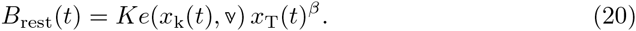

Substituting it in (10), Eq. (20) captures the notion that energy extraction gives the individual energy that it can use to grow or maintain its different tissues.

### Closing the model

The expression for the resting metabolic rate [Eq. (20)] closes the model from a metabolic point of view, since after substituting Eq. (20) in Eq. (10) [and using Eqs. (11) and (12)], the ontogenetic dynamics of the brain, reproductive, and somatic tissue mass, *x*_b_(*t*), *x*_r_(*t*), and *x*_s_(*t*), and of the number of skills, *x*_k_(*t*), are expressed in terms of such state variables, of empirically estimable parameters, and on the evolving traits (mutant 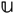 and resident 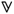). To close the model from an evolutionary perspective and compute an optimal allocation strategy, we need expressions for how the state variables relate to the vital rates [*l*(*t*) and *m*(*t*)] in Eq. (2) and expressions for the energy extraction efficiency 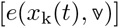. A large number of different settings can be conceived with the model so far, both for the vital rates and energy extraction efficiency. We focus on an application aiming at modeling human brain evolution from the baseline setting “me-against-nature” to be compared with future elaborations of the model.

#### Vital rates

For simplicity, we consider that the mortality rate *μ* of an individual is independent of age and of the evolving traits, and so

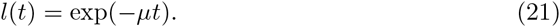

We also assume that density-dependent regulation acts on fecundity (e.g., through lottery competition) so that the effective fecundity *m*(*t*) is proportional to fecundity *f* (*t*), defined as the rate of offspring production at age t without density dependence (e.g., [50, 53, 61]). That is, we let

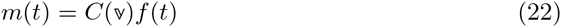

where 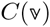 is a proportionality factor that depends on population size which ultimately depends on the resident strategy 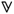.

We obtain a measure of fecundity *f*(*t*) with an analogous reasoning to that used for the learning rate of skills. In particular, we assume that some of the metabolic rate of the reproductive tissue is due to offspring production and maintenance. Denote by *M*_repr_(*t*) the metabolic rate of the reproductive tissue at age *t* (i.e., the heat released by the reproductive tissue per unit time with the individual at rest). From energy conservation and Eqs. (3) and (4), the reproductive metabolic rate must satisfy

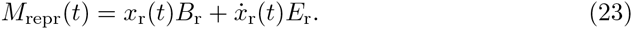

Let *s*_r_ be the fraction of the reproductive metabolic rate allocated to offspring production and maintenance, which we assume constant for simplicity. Let *x_o_*(*t*) be the number of offspring the individual has at age *t*. Suppose that reproductive tissue releases an amount of heat *E_o_* for the production of an average offspring (fecundity cost). Similarly, assume that the reproductive tissue releases an amount of heat *B_o_* per unit time for maintaining an average offspring (physiological cost of maternal care; e.g., due to lactation). Hence, from energy conservation, the rate of heat release by the reproductive tissue due to offspring production and maintenance must equal the reproductive metabolic rate allocated to offspring production and maintenance:

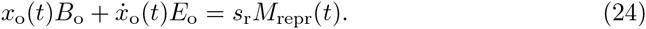

Rearranging, we obtain the model’s fourth key equation specifying fecundity:

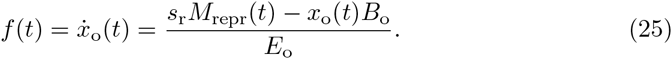

The first term in the numerator of (25) gives the heat released due to energetic input for reproduction whereas the second term gives the heat released for physiological parental care.

Eq. (25) can be simplified as follows. If the reproductive tissue is defined narrowly enough (e.g., as preovulatory ovarian follicles) so that it is not involved in offspring maintenance, the physiological costs of maternal care incurred by the reproductive tissue are essentially null (i.e., *B*_o_ ≈ 0; they are, however, included in the maintenance costs *B*_s_ of the somatic tissue as we defined it above). With this definition of reproductive tissue, we take body mass at age *t* as *x*_T_(*t*) = *x*_b_(*t*) + *x*_r_(*t*) + *x*_s_(*t*). If additionally, reproductive tissue maintenance is much more expensive than production (i.e., *B*_r_ ≫ *E*_r_, which holds with our estimated parameters for humans; Table S2), fecundity can be approximated as

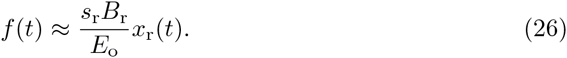

For the results reported below, the approximation (26) is accurate and yields no detectable difference in the predicted adult brain and body mass [Supporting Information (SI) §7].

Using Eq. (22) and (26), effective fecundity becomes

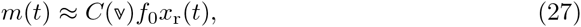

where *f*_0_ = *s*_r_*B*_r_/*E_o_*. We assume that non-physiological costs of (allo)parental care are included in 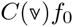. Effective fecundity is then proportional to the mass of reproductive tissue, which is consistent with medical approaches to predict fecundity in women in terms of ovarian follicle count [62,63].

#### Energy acquisition

We now model energy acquisition. We assume that energy extraction at age t is done exclusively by overcoming a challenge posed by the non-social environment (e.g., gathering food or lighting a fire) and that the individual engages alone (but possibly with caregivers’ help) in overcoming such a challenge (“me against nature”). This setting implies that the energy extraction efficiency,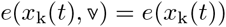 is independent of the resident strategy.

We treat the me-against-nature setting as a contest against the environment. We thus let energy extraction efficiency *e*(*x*_k_(*t*)) take the form of a contest success function [64,65]:

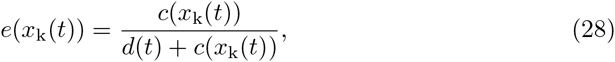

which depends on two terms. First, energy extraction efficiency depends on the difficulty of the challenge at age *t*, measured by *d*(*t*). The higher *d*(*t*), the more challenging energy extraction is and the more energy-extraction skills the individual must have to obtain resources. We let *d*(*t*) = α – *φ(t)*, where *α* is the environmental difficulty and *φ(t)* is the facilitation of the challenge due to (allo)parental care. We let this facilitation be an exponentially decreasing function with age, *φ(t)* = *φ*_0_ exp(–φ_r_*t*), and for simplicity we ignore the increased resting metabolic rate caused by gestation and lactation [66].

Second, energy extraction efficiency depends on the individual’s competence, denoted by *c*(*x*_k_(*t*)). We consider two cases that are standard in contest models: (1) a power function *c*(*x*_k_(*t*)) = (*x*_k_(*t*))^*γ*^, so energy extraction efficiency *e*(*x*_k_(*t*)) is a contest success function in ratio form (power competence); and (2) an exponential function *c*(*x*_k_(*t*)) = (exp(*x*_k_(*t*)))^*γ*^ so energy extraction efficiency is in difference form (exponential competence) [64,65]. In both cases, the parameter *γ* describes the effectiveness of skills at energy extraction. Thus, with *γ* =0, skills are ineffective while with increasing *γ* fewer skills are needed to extract energy. In general, competence *c*(*x*_k_(*t*)) represents features of the individual (e.g., how increasing skill changes efficiency in information processing by the brain), and of the environment (e.g., how adding the skill of caching nuts to that of cracking nuts changes energy extraction efficiency). For a given skill effectiveness (*γ*), exponential competence assumes a steeper increase in competence with increasing skill number than power competence.

#### Model summary and solution implementation

On substituting Eqs. (21) and (27) into Eq. (2) along with Eq. (28) into Eq. (20), the model is closed and can be used to determine uninvadable allocation strategies and the resulting equilibrium growth patterns. From our simplifying assumptions, determining uninvadable allocation strategies reduces to an optimal control problem. We obtain locally uninvadable allocation strategies using optimal control methodology (e.g., [67]), both by a “direct” approach with the software GPOPS [68] for numerical approximations and by an “indirect” approach using Pontryagin’s maximum principle for analytical results (SI §1–4). The model depends on 22 parameters which measure (P1) tissue mass in the newborn (*x*_*i*0_ for *i* ∈ {*b, r, s*}), (P2) tissue metabolism (*K, β, B_i_* and *E_i_* for *i* ∈ {*b, r, s*}), (P3) demography (*f*_0_, *μ*, and *T*), (P4) skill of the newborn (*x*_k0_), (P5) skill metabolism (*s*_k_, *B*_k_ and *E*_k_), (P6) (allo)parental care (*φ*_0_ and *φ*_r_), and (P7) contest success (*α* and *γ*). From their definitions, the parameters are measured in units of mass, energy, time, and skill. The parameter *f*_0_ only displaces the objective vertically and thus has no effect on the uninvadable allocation strategies. The parameter *T* is taken finite for numerical implementation and set as the observed age of menopause.

We use published data for human females to estimate 13 parameters that affect the uninvadable allocation strategies (P1-P3) (SI §5,6; Table S2). These parameters include the brain and body metabolic costs, and with these parameters fixed, the model can only generate a vastly narrower set of outcomes. These parameters are in units of mass, energy, and time which we measure in kg, MJ (megajoules), and years, respectively. The remaining 8 parameters that affect the uninvadable allocation strategies (P4-P7) are less easily estimated from available data, so we identify by trial-and-error benchmark values that yield a model output in agreement with observed ontogenetic body and brain mass data for modern human females. The benchmark parameter values are different with power (Table S3) and exponential (Table S4) competence. The benchmark parameter values involve skill units, and as we do not estimate them from empirical data, we measure skill in arbitrary units. We first present the numerical results for the two sets of benchmark parameter values and then the results when deviating from them (see SI for analytical results and computer code).

## Results

### Predicted life history stages: childhood, adolescence, and adulthood

The optimal strategy we obtain divides the individual’s lifespan in three broad stages: (1) a “childhood” stage, defined as the stage lasting from birth to *t*_m_ years of age (age at maturity) and during which allocation to growth of reproductive tissue is zero; (2) an “adolescence” stage, defined as the stage lasting from *t*_m_ to *t*_a_ years of age (age at adulthood) and during which there is simultaneous allocation to growth of somatic and reproductive tissue; and (3) an “adulthood” stage, defined as the stage lasting from ta to the end of the individual’s reproductive career and during which all growth allocation is to reproductive tissue (Fig. 2A). These life stages are obtained with either power or exponential competence (Fig. 2A,E). Note that the ages at maturity *t*_m_ and adulthood ta (switching times) are not parameters but an output of the model.

**Fig. 2.**
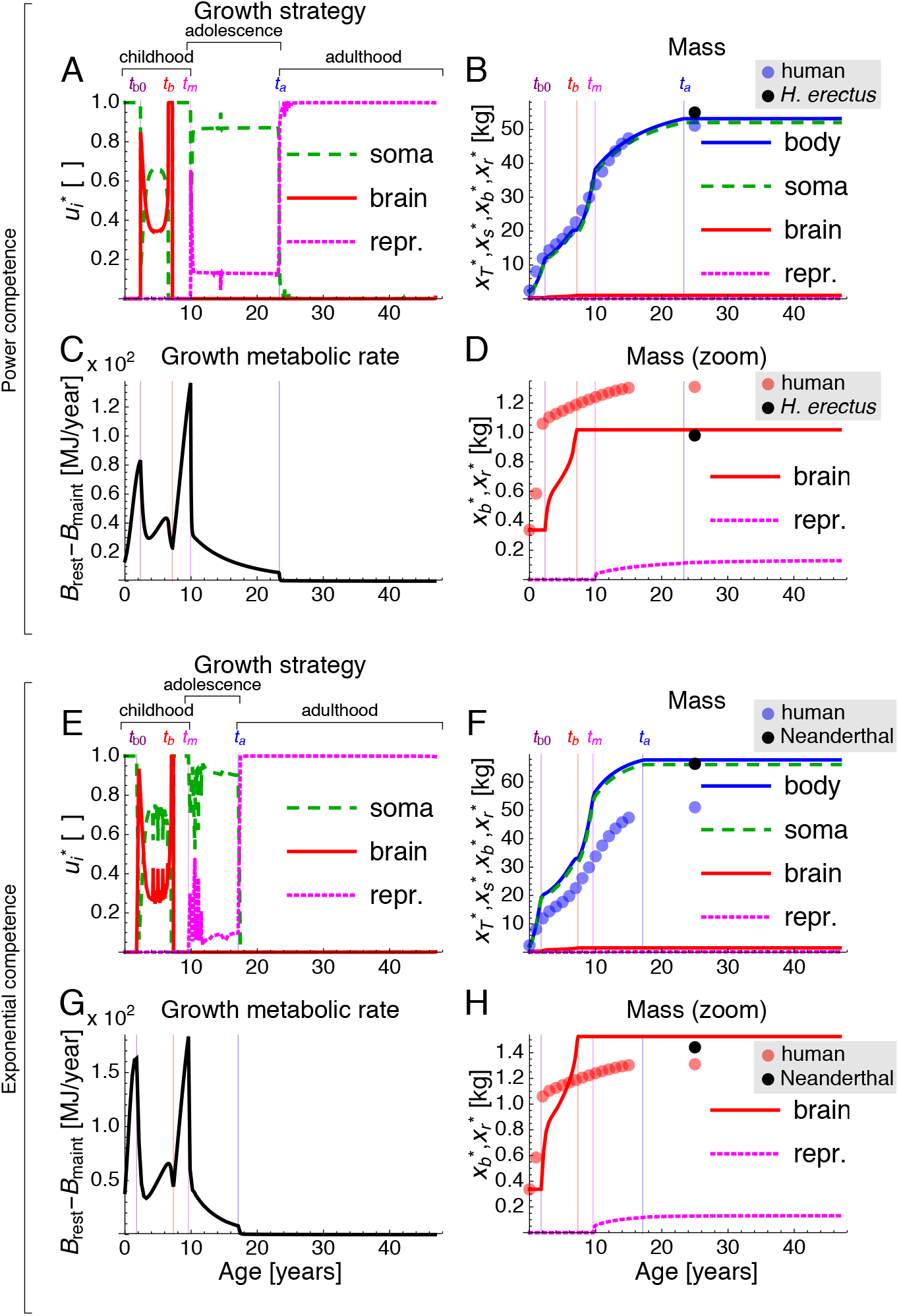
Uninvadable growth strategy 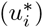 and the resulting growth patterns 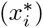 under a me-against-nature setting. Lines are model’s results and circles are observed values in human females. Results with (A-D) power and (E-H) exponential competence. (A,E) Uninvadable growth strategy vs. age. (C,G) Resulting growth metabolic rate vs. age. (B,F) Resulting body and tissue mass vs. age. (D,H) Resulting brain and reproductive mass vs. age. Lines and circles with the same color are respectively the model’s prediction and the observed values in modern human females [69]. Black circles are the observed (B,F) adult female body mass and (D,H) adult sex-averaged brain mass, either for late *H. erectus* [70] or Neanderthals [71,72]. Jitter in the growth strategy (A,E) is due to negligible numerical error (Fig. S2).

The obtained childhood stage, which is the only stage where there is brain growth, is further subdivided in three periods: (1a) “ante childhood”, defined here as the earliest childhood period with pure allocation to somatic growth; (1b) “childhood proper”, defined here as the childhood period where there is simultaneous allocation to somatic and brain growth; and (1c) “preadolescence”, defined here as the latest childhood period of pure somatic growth. Hence, brain growth occurs exclusively during “childhood proper”. The occurrence of an “ante childhood” without brain growth disagrees with observation in humans. Two possible and particularly relevant reasons for this discrepancy may be either the absence of social interactions in this setting of the model, or the approximation of resting metabolic rate by a power law (20) which underestimates resting metabolic rate (and thus growth metabolic rate) during ante childhood (Fig. S3). The period we refer to here as childhood proper then lasts from the obtained age *t*_b0_ of brain growth onset to the obtained age *t*_b_ of brain growth arrest (these switching times are also an output rather than parameters of the model; Fig. 2A).

With the exception of the age of brain growth onset, the predicted timing of childhood, adolescence, and adulthood closely follows that observed in humans with competence being either a power or an exponential function of skill number, given their respective benchmark parameter values (Table 1). Recall that measurement units (i.e., years, kg, and MJ), excepting skill units, are not arbitrary as they result from the units of the parameter values estimated from empirical data (Table S2). Hence, while using realistic metabolic costs of brain and body, the model can correctly predict major stages of human life history with accurate timing, with the possible exception of brain growth allocation during ante childhood (Table 1).

**Table 1.**
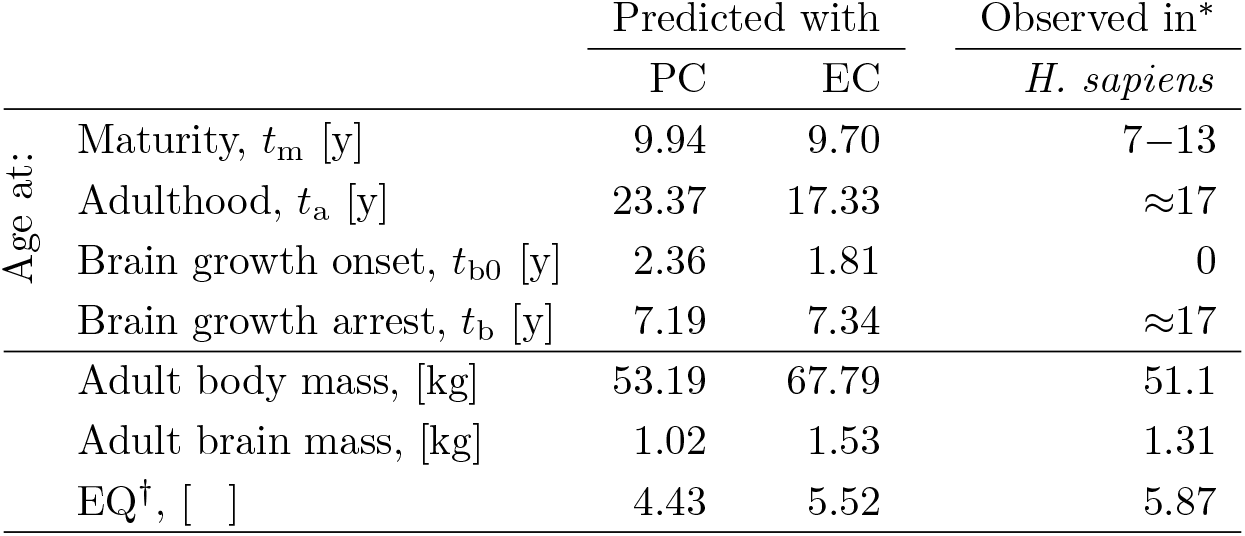
Predictions for life history timing and adult brain and body mass with a me-against-nature setting. Switching times and adult values resulting with competence as a power or exponential function (PC and EC) for the results in Fig. 2. * Observed values in human females: age at maturity [73], adulthood [74], brain growth onset and arrest [69], adult body mass [69], and adult brain mass [69]. †Encephalization quotient, calculated as EQ = *x*_b_(*t*_a_)/ [11.22 × 10^−3^*x*_T_(*t*_a_)^0.76^] (mass in kg) [75].

### Body and brain mass through ontogeny

The optimal growth strategy generates the following predicted body and brain mass throughout ontogeny. For total body mass, there is fast growth during ante childhood, followed by slow growth during childhood proper, a growth spurt during preadolescence, slow growth during adolescence, and no growth during adulthood, each of which closely follows the observed growth pattern in humans (Fig. 2B). The slow growth during childhood proper results from the simultaneous allocation to somatic and brain growth and from the decreasing growth metabolic rate due to the increasing energetic costs of brain maintenance (Fig. 2C). The growth spurt during preadolescence arises because (1) all growth metabolic rate is allocated to inexpensive somatic growth, and (2) growth metabolic rate increases due to increased metabolic rate caused by increasing, inexpensive-to-maintain somatic mass (Fig. 2C). The slow growth during adolescence is due to simultaneous somatic and reproductive growth, and to the elevated costs of reproductive tissue maintenance (Fig. 2C). These growth patterns result in two major peaks in growth metabolic rate (Fig. 2C). While the first peak in growth metabolic rate is made possible by (allo)parental care, the second peak is made possible by the individual’s own skills (Fig. S8D). After the onset of adulthood at *t*_a_, growth metabolic rate is virtually depleted and allocation to growth has essentially no effect on tissue growth (Fig. 2C).

Whereas predicted body growth patterns are qualitatively similar with either power or exponential competence, they differ quantitatively (Fig. 2B,F). With power competence, the predicted body mass is nearly identical to that observed in human females throughout life (Fig. 2B). In contrast, with exponential competence, the predicted body mass is larger throughout life than that of human females (Fig. 2F). Our exploration of the parameter space indicates that the larger body mass with exponential competence relative to power competence is robust to parameter change (Figs. 4A-C, S16A-C, S17A-F).

Regarding brain mass, the model predicts it to have the following growth pattern. During ante childhood, brain mass remains static, in contrast to the observed pattern (Fig. 2D). During childhood proper, brain mass initially grows quickly, then it slows down slightly, and finally grows quickly again before brain growth arrest at the onset of preadolescence (Fig. 2D). Predicted brain growth is thus delayed by the obtained ante-childhood period relative to the observed brain growth in humans (Fig. 2D). As previously stated, such brain growth delay may be a result of the absence of social interactions in this model setting, or an inaccuracy arising from the underestimation of resting metabolic rate during ante childhood by the power law of body mass.

Predicted brain growth patterns are also qualitatively similar but quantitatively different with power and exponential competence (Fig. 2D,H). Adult brain mass is predicted to be larger with competence as an exponential rather than as a power function (Fig. 2D,H). As for body mass, our exploration of the parameter space indicates that the larger brain mass with exponential competence is robust to parameter change (Figs. 4A-C, S16A-C, S17A-F). Moreover, the encephalization quotient (EQ, which is the ratio of observed adult brain mass over expected adult brain mass for a given body mass) is also larger with exponential competence for the benchmark parameter values (Table 1). For illustration, with competence as a power function, the predicted adult body and brain mass approach those observed in late *H. erectus* (Fig. 2B,D). In contrast, with competence as an exponential function, the predicted adult body and brain mass approach those of Neanderthals (Fig. 2F,H). The larger EQ with exponential competence is also robust to parameter change (Figs. 4D-F, S16D-F, S17G-L).

### Skills through ontogeny

The obtained optimal growth strategy predicts the following patterns for energy-extraction skills throughout ontogeny. Under the same parameter values as in Fig. 2, the individual gains most skills during childhood and adolescence, skill number continues to increase after brain growth arrest, and skill number plateaus in adulthood (Fig. 3). That is, skill growth is “determinate”, in agreement with empirical observations (Fig. 3). Yet, if memory cost *B*_k_ is substantially lower, skill number can continue to increase throughout life (i.e., skill growth is then “indeterminate”; Fig. S9E) [see Eq. (14)]. Nevertheless, in that case, the agreement between predicted and observed body and brain mass throughout ontogeny is substantially reduced (Fig. S9B,C).

**Fig. 3.**
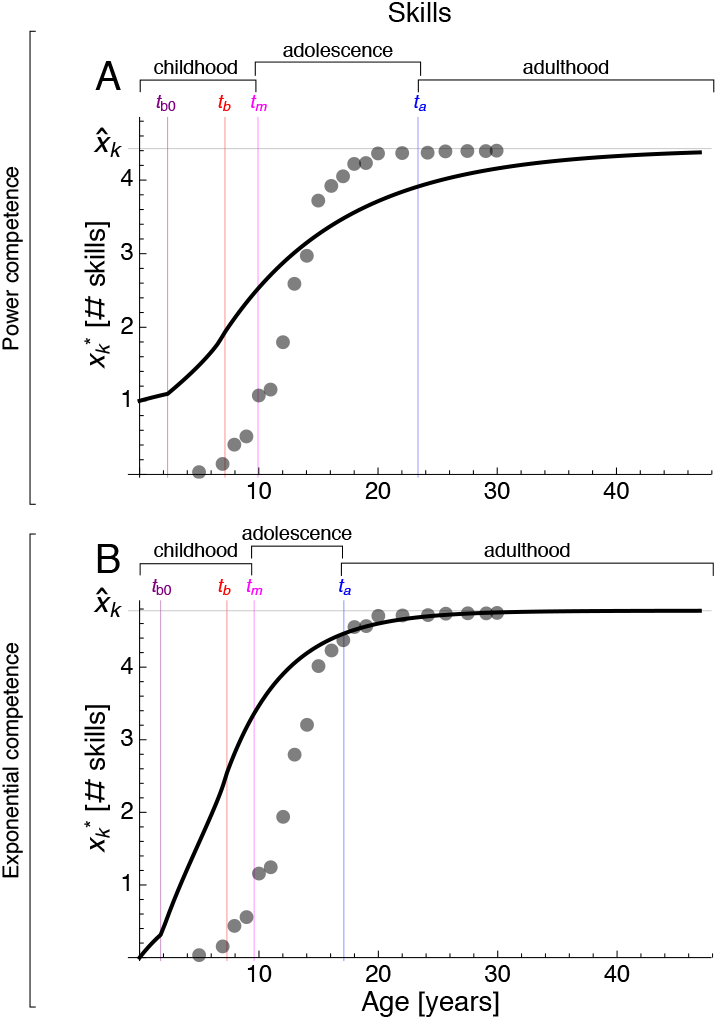
Predicted skill ontogeny plateaus within the individual’s lifespan. Lines are the predicted number of skills vs. age with power (A) and exponential (B) competence for the results in Fig 2. Circles are the observed cumulative distribution of self-reported acquisition ages of food production skills in female Tsimane horticulturalists [76] multiplied by our 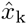. However, note that the observed skills in Tsimane include socially learned skills which we do not consider explicitly in the model.

When skill growth is determinate, the model predicts adult skill number to be proportional to adult brain mass. In particular, with determinate skill growth, the number of skills that is asymptotically achieved [from Eq. (14) setting 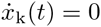 and 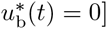] is

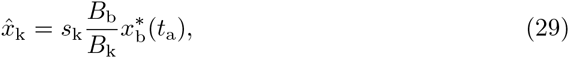

where 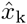 is the asymptotic skill number, 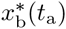 is the adult brain mass, *s*_k_ is the fraction of brain metabolic rate allocated to energy-extraction skills, and *B*_b_ is the brain mass-specific maintenance cost. The requirement for skill growth to be determinate is that the brain metabolic rate allocated to skills [*s*_k_M_brain_(*t*)] becomes saturated with skill maintenance [*x*_k_(*t*)*B*_k_] within the individual’s life [Eq. (14)]. Hence, adult skill number is proportional to adult brain mass in the model because of saturation with skill maintenance of the brain metabolic rate allocated to skills and because adult brain metabolic rate is found to be proportional to adult brain mass [setting 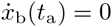 in Eq. (12) yields *M*_brain_(*t*_a_) = *x*_b_(*t*_a_)*B*_b_]. Weak correlations between cognitive ability and brain mass have been identified across taxa including humans [5,77–80]. Since skills are here broadly understood to include cognitive abilities (provided parameters are suitably reinterpreted), this result offers an explanation for these correlations in terms of saturation of brain metabolic rate with skill maintenance (memory).

We now vary parameter values to assess what factors favor a large brain at adulthood in a me-against-nature setting.

### A large brain is favored by intermediate environmental difficulty, moderate skill effectiveness, and costly memory

A larger adult brain mass is favored by an increasingly challenging environment [increasing α; Eq. (28)], but is *disfavored* by an exceedingly challenging environment (Fig. 4A). Environmental difficulty favors a larger brain because more skills are needed for energy extraction [Eq. (28)], and from Eq. (14) more skills can be gained by increasing brain metabolic rate in turn by increasing brain mass. Thus, a large brain is favored to energetically support skill growth in a challenging environment. However, with exceedingly challenging environments, the individual is favored to reproduce early without substantial body or brain growth because it fails to gain enough skills to maintain its body mass as (allo)parental care decreases with age (Fig. S13).

**Fig. 4.**
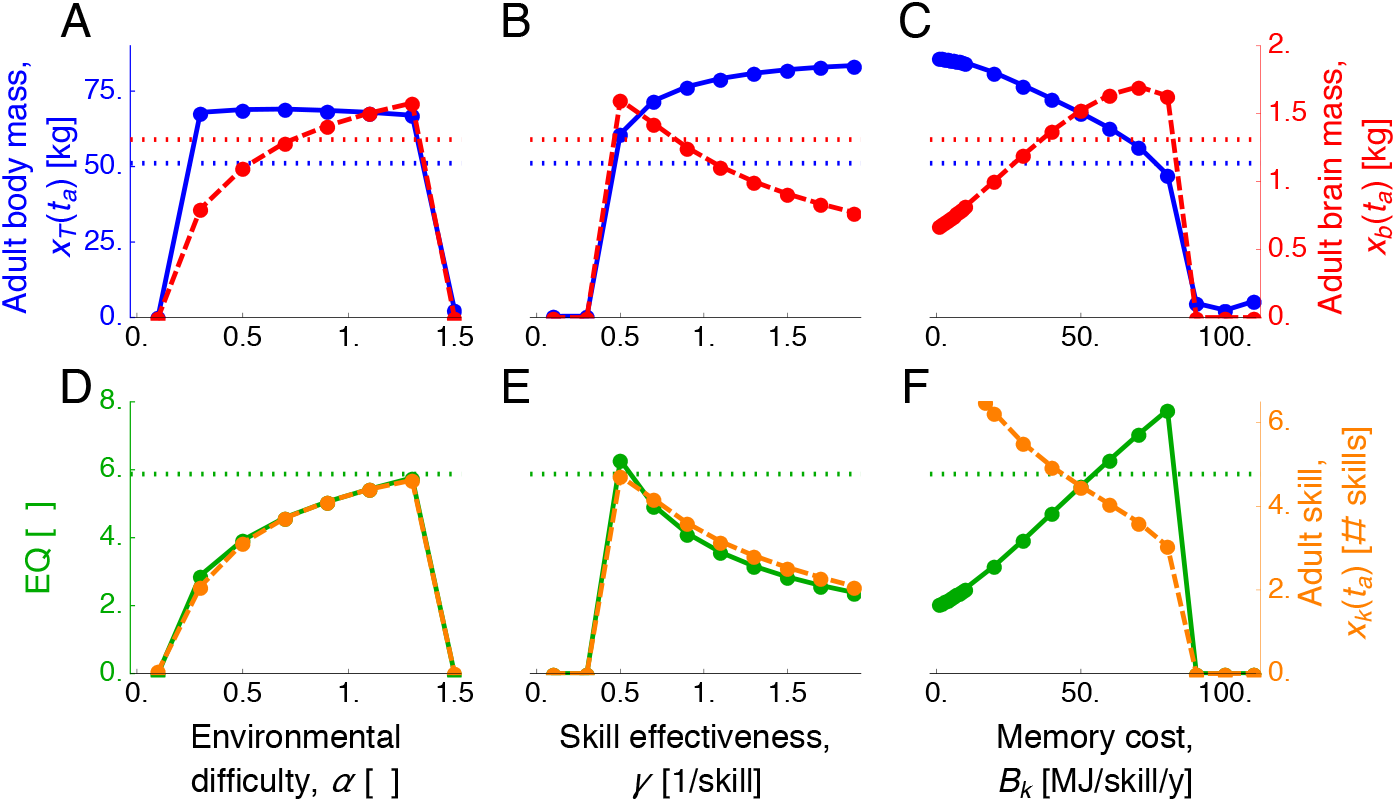
Large adult brain mass and EQ are favored by environmental difficulty, moderate skill effectiveness, and costly memory. Plots are the predicted adult body and brain mass, EQ, and skill vs. parameter values with exponential competence. A-C show adult body mass (blue) and adult brain mass (red). D-F show adult EQ (green) and skill (orange). Vertical axes are in different scales. Dashed horizontal lines are the observed values in human females [69].

A larger adult brain is favored by moderately effective skills. When skills are ineffective at energy extraction [*γ* → 0; Eq. (28)], the brain entails little fitness benefit and fails to grow in which case the individual also reproduces without substantially growing (Fig. 4B). When skill effectiveness (*γ*) crosses a threshold value, the fitness effect of brain becomes large enough that the brain becomes favored to grow. Yet, as skill effectiveness increases further and thus fewer skills are needed for energy extraction, a smaller brain supports enough skill growth, so the optimal adult brain mass *decreases* with skill effectiveness (Fig. 4B). Hence, adult brain mass is largest with moderately effective skills.

A larger brain is also favored by skills that are increasingly expensive for the brain to maintain (costly memory, increasing *B*_k_), but exceedingly costly memory prevents body and brain growth (Fig. 4C). Costly memory favors a large brain because then a larger brain mass is required to energetically support skill growth [Eq. (14)]. If memory is exceedingly costly, skills fail to grow and energy extraction is unsuccessful, causing the individual to reproduce without substantial growth (Fig. 4C).

### Factors favoring a large EQ and high skill

A large EQ and high adult skill number are generally favored by the same factors that favor a large adult brain. However, the memory cost has a particularly strong effect favoring a large EQ because it simultaneously favors increased brain and reduced body mass (Fig. 4C,F). In contrast to its effect on EQ, increasing memory cost *disfavors* a high adult skill number (Fig. 4F). That is, a higher EQ attained by increasing memory costs is accompained by a *decrease* in skill number (Fig. 4C,F). The factors that favor a large brain, large EQ, and high skill are similar with either power or exponential competence (Fig. 4 and Figs. S16, S17). Importantly, although with the estimated parameter values the me-against-nature setting can recover human growth patterns yielding adult body and brain mass of ancient humans, our exploration of the parameters that were not estimated from data suggests that the me-against-nature setting cannot recover human growth patterns yielding adult body and brain mass of *modern* humans.

## Discussion

By combining elements of life history and metabolic theories, we formulated a metabolically explicit mathematical model for brain life history evolution that yields testable quantitative predictions from predefined settings. We analyzed the model for a me-against-nature setting where individuals have no social interactions except possibly with caregivers, but the model can be implemented to study brain evolution more generally. Our results for the me-against-nature case show that this setting can be sufficient to generate major human life history stages as well as adult brain and body mass of ancient human scale, all without social interactions or evolutionary arms races in cognition triggered by social conflict. Overall, we find that in the model the brain is favored to grow to energetically support skill growth, and thus a larger brain is favored when (1) competence at energy extraction has a steep dependence on skill number, (2) many skills are needed for energy extraction due to environmental difficulty and moderate skill effectiveness, and (3) skills are expensive for the brain to maintain but are still necessary for energy extraction.

The model correctly divides the individual’s lifespan into childhood, adolescence, and adulthood. The model also rightly predicts brain growth to occur only during childhood, although there is a delay in the predicted brain growth which may be due to the absence of social interactions or an underestimation of resting metabolic rate early in life by its power law approximation. Additionally, the predicted childhood stage finishes with a growth spurt, as observed in human preadolescence. The model also recovers an adolescence stage with simultaneous allocation to growth and reproduction, which has previously been difficult to replicate with life history models [28]. While the timing of these predicted life stages depends on the magnitude of parameter values, their relative sequence is likely to depend on the relative magnitude of metabolic costs of maintenance and production of the different tissues (i.e., on whether (1) *B_i_* < *B_j_* and (2) *E_i_* < *E_j_* for *i, j* ∈ {*b, r, s*}). Empirically guided refinement of both parameter values and the shape of energy extraction efficiency is expected to allow for increasingly accurate predictions [81]. Similarly, empirical data for non-human taxa should allow determining the model’s ability to predict diverse life histories and brain growth patterns [82].

The model also offers an explanation for observed inter- and intraspecific correlations between adult cognitive ability and brain mass across taxa including humans [5,77–80]. The explanation is the saturation with memory costs of the brain metabolic rate allocated to skills during the individual’s lifespan [Eq. (29)]. The proportionality arises because the adult brain metabolic rate is found to be proportional to brain mass. This explanation follows from a general equation for the learning rate of skills [Eq. (14)] that is based on metabolic considerations [34] without making assumptions about skill function; yet, this equation assumes that the fraction of brain metabolic rate allocated to the skills of interest (*s*_k_) is independent of brain mass (and similarly for *B*_b_ and *B*_k_). The model further predicts that additional variation in correlations between cognitive ability and brain mass can be explained by variation in maintenance costs of brain and skill, and by variation in brain metabolic rate allocation to skill [Eq. (29)]. However, the model indicates that adult skill number and brain mass need not be correlated since saturation with skill maintenance of the brain metabolic rate allocated to skills may not occur during the individual’s lifespan, for example if memory is inexpensive, so skill number increases throughout life (Fig. S9E).

Predicted adult brain mass and skill have non-monotonic relationships with their predictor variables (Figs. 4, S16, S17). Consequently, conflicting inferences can be drawn if predictor variables are evaluated only on their low or high ends. For instance, increasingly challenging environments favor large brains up to a point, so that exceedingly challenging environments disfavor large brains. Thus, on the low end of environmental difficulty, the prediction that increasingly challenging environments favor large brains is consistent with ecological challenge hypotheses [21,38]; yet, on the high end of environmental difficulty, the prediction that increasingly challenging environments disfavor large brains is consistent with constraint hypotheses according to which facilitation of environmental challenge favors larger brains [21,83–85]. Counter-intuitively on first encounter, the finding that moderately effective skills are most conducive to a large brain and high skill is a consequence of the need of more skills when their effectiveness decreases (Fig. 4B). Regarding memory cost, the strong effect of memory cost on favoring a high EQ at first glance suggests that a larger EQ than the observed in humans is possible if memory were costlier (see dashed lines in Fig. 4E). However, such larger memory costs cause a substantial delay in body and brain growth, and the resulting growth patterns are inconsistent with those of humans (Figs. S10-S12).

Although our model does not include numerous details relevant to humans including social interactions and social learning, our results are relevant for a set of hypotheses for human-brain evolution. In particular, food processing (e.g., mechanically with stone tools or by cooking) has previously been advanced as a determinant factor in human-brain evolution as it increases energy and nutrient availability from otherwise relatively inaccessible sources [86,87]. Evidence of human fire control has been inconclusive for early dates (1.5 mya, associated with early H. *erectus* in South Africa), while being more secure for more recent dates (800 kya, associated with *H. erectus* in Israel) and abundant for yet more recent times (130 kya, associated with Neanderthals and *H. sapiens* throughout the Old World) [88,89]. Evidence of fire deep inside a South African cave associated to *H. erectus* has been identified for sediments dated to 1 mya [90]. Regarding mechanical processing, “many of the oldest stone tools bear traces of being used to slice meat” (1.5 mya in Kenya; [87,91]) and experimental evidence shows that meat slicing and vegetable pounding substantially reduce chewing effort [87]. Food processing relates to our results not only in that it can primarily constitute a me-against-nature setting, but also in that it may help satisfy at least two of the three key conditions identified for large-brain evolution listed in the first paragraph of the Discussion. First, a shift in food-processing technology (e.g., from primarily mechanical to cooking) could create a steeper relationship between energy-extraction skills and competence by substantially facilitating energy extraction (relating to condition 1). Second, food processing (e.g., by building the required tools or lighting a fire) is a challenging feat to learn and may often fail (relating to condition 2). Yet, there are scant data allowing to judge the metabolic expense for the brain to maintain tool-making or fire-control skills (condition 3). Our results are thus consistent with the hypothesis of food processing as being a key factor in human brain expansion.

In sum, the model identifies various drivers of large-brain evolution, in particular steep competence with respect to skill, intermediate environmental difficulty, moderate skill effectiveness, and costly memory. As we did not consider social interactions, our application of the model cannot refute or support social brain hypotheses. However, application of our model to the social realm should allow for assessments of social hypotheses.

## Supporting Information

**Supporting Information. SI.** Analytical results, parameter estimation, and supplementary numerical results.

**Supporting Data. SD.** MATLAB computer code for solutions using GPOPS.

## Acknowledgments

We thank Tadeusz J. Kawecki for helpful discussion.

## Author Contributions

MGF and LL derived the model. MGF obtained the results. TF contributed analytic tools. MGF and LL wrote the paper.

